# Assessing the efficiency and the side effects of atrazine-degrading biocomposites amended to atrazine-contaminated soil

**DOI:** 10.1101/2025.04.08.647713

**Authors:** Sakineh Abbasi, Marion Devers-Lamrani, Fabrice Martin-Laurent, Caroline Michel, Sana Romdhane, Nadine Rouard, Aymé Spor

## Abstract

Even decades after being banned in Europe, atrazine and its main metabolites can still be found in soils. While bioaugmentation using pesticide-degrading bacteria is already employed as a strategy for remediating polluted soils, there is still a need to improve its efficiency. Therefore, investigating the application of carrier materials to deliver and stabilize pesticide-degrading microorganisms *in situ* emerges as an interesting approach for further exploration. Here, we generated atrazine-degrading biocomposites by cultivating either a single strain or a 4-species bacterial consortium as biofilms on zeolite, which serves as the carrier material. Using a microcosm experiment, we then evaluated their efficiency to mineralize ^14^C-atrazine in an agricultural soil comparing to free-living cells, and assessed the side effects of the two inoculation methods on the native soil bacterial community using 16S rDNA amplicon sequencing. We showed that, right after inoculation, the atrazine mineralization potential of the free-living cells was higher than that of the biocomposites. However, microcosms inoculated with the biocomposites displayed significantly higher atrazine mineralization potential than the ones inoculated with free-living cells after 15 and 45 days of incubation, not only indicating a higher efficiency but also a better stability in the soil environment, further confirmed by qPCR of the *atz* genes. We also showed that the inoculation of free-living cells and biocomposites differently influences the diversity and composition of the native microbial community, and that these effects are modulated by the scenario of atrazine contamination during soil inoculation. Altogether, our results provide a thorough evaluation of the efficiency and the ecotoxicological impact of atrazine-degrading biocomposites in soil.

## INTRODUCTION

Atrazine is one of the most popular and widely used herbicides in the world because of its efficiency and low cost in controlling broadleaf and grassy weeds (Rostami et al., 2021). Its use was banned in Europe in 2003 notably because of ubiquitous and unpreventable water resources contamination. Atrazine and its degradation products can persist in soils for decades after applications, and concerns are growing regarding its potential toxicity for animal and human health (Rostami et al., 2021). Hence, there is a strong need to find solutions to improve the efficiency of removing long-lasting atrazine contaminations from the environment.

Several studies have revealed that bioremediation strategies increase the degradation of atrazine in contaminated soils (Dominguez-Garay et al., 2016; Sanchez et al., 2019; Chen et al., 2019). The application of atrazine-degrading bacteria, either as a single species or in consortium, can enhance atrazine removal from the soil (Struthers et al., 1988). However, their success relies on both the stability of atrazine-mineralization and the survival rate of inoculated strains (Zhu et al., 2019; Zhao et al., 2018). Microbial consortia might have an advantage over isolated strains due to synergistic interactions and/or labour sharing between diverse species and are often characterized by a higher degradation efficiency of complex compounds (Zhang and Zhang, 2022). For example, in a previous study, we showed that the establishment of coexistence between species, through atrazine-degrading gene loss, allowed the members of a bacterial consortium to efficiently improve the degradation of atrazine (Billet et al., 2019).

The establishment of microbial inoculants in soil is challenged by both abiotic and biotic factors. The soil physico-chemical properties, as well as the presence of an already established autochthonous soil microbiota often act in concert to prevent the invasion of alloctonous microbial inoculants (Borges et al., 2019). Therefore, innovative technologies need to be explored to improve the stability and efficiency of inoculants in contaminated soils. An effective solution involves the immobilization of bacteria on zeolite as a microbial carrier, thereby offering protection and improving their stability in soils (Gorodylova et al., 2021). Natural zeolite is a commonly occurring sedimentary deposit that is microporous, crystalline, negatively charged, inexpensive, and non-toxic (Zheng et al. 2001). The generation of biofilm on the surface of the carrier material is dependent on the interaction between bacteria and support, which leads to microbial adhesion and colonization of the support surface (Zheng et al., 2001; Jiang et al., 2007). Some properties of natural or modified zeolite materials such as their adsorption capacities and structural stabilities could enable them to be used as a support to prevent losing free-living cells during bioremediation (Mohsin et al., 2023; Kuldeyev et al., 2023). However, it is important to explore not only the benefits of these supports on the stability and efficiency of degrading inoculants in removing atrazine residues but also their potential side effects on autochtonous soil microbial communities (Jia et al., 2021).

Here, we compared the efficiency and stability of a single species and a four-species consortium-based biocomposites to remediate a simulated atrazine contamination in soil. We also investigated whether these biocomposites have side effects on native microbial communities. We hypothesized that *i*) the consortium-based biocomposite would be more efficient than the single species-based biocomposite in soil, *ii*) the biocomposite atrazine-degrading inoculants would be more stable and efficient in removing atrazine contamination in soil than free-living cells inoculant, *iii*) the biocomposites would have a stronger impact than the free-living cells on the structure of the autochthonous microbial community.

## MATERIALS AND METHODS

### Strains and media

The pure bacterial strain used in this study is *Pseudomonas* sp. ADP3. It derives from *Pseudomonas* sp. ADP (de Souza *et al*., 1996) and harbours the *atzABCDEF* genes. The bacterial consortia used was made of *Variovorax* sp. 38R (*atzA*), *Chelatobacter* sp. *SR38* (*atzCtrzD*)*, Pseudomonas* sp. ADPe (*atzDEF*), and *Arthrobacter* sp. TES (*trzNatzBC*) (Billet et al., 2019 & 2021). The pure strain and the consortia were cultivated in MSA medium, a mineral salt liquid medium containing 4 mM of sodium citrate as a carbon source and 0.3 mM of atrazine as the sole nitrogen source (Mandelbaum et al., 1993). For enumeration, TY solid medium was used (bactotryptone 5 g L^-1^, yeast extract 3 g L^-1^, CaCl_2_ 0.6 mM).

### Soil and mineral support

Soil samples were collected from the top horizon layer (0-20 cm) of the Epoisses INRAE experimental farm in France (47° 30’ 22.1832’’ N, 4° 10’ 26.4648’’ E). Soil properties are as follows: 51.9% silt, 41.9% clay, 6.2% sand, 15.5 g/kg of organic carbon, 1.4 g/kg of nitrogen content, and pH 7.2. Soil samples were sieved at 4 mm and stored at 4°C temperature less than two months before use.

Natural mineral zeolite used in this study was purchased from Saint-Malo, France (z-SM) and prepared as already described (Gorodylova et al., 2021). The sample was ground using a hummer crusher and its particle size was adjusted in the range 0.2–1.25 mm using laboratory sieves (diameter: 20 cm; openings: 0.2, 0.315, 0.4, 0.5, 0.63, 0.8, 1 and 1.25 mm). In order to remove dust, wood and plastic contaminants, the sample was washed by floatation in deionised water, filtered and dried at 60 °C for 24 h. It was then further sterilized by autoclaving.

### Biocomposite preparation

To prepare the biocomposites, a preculture of the pure strain and of the consortia was performed from a stock frozen cell suspension inoculated in MSA at 28°C under agitation (120 rpm). After 4 days of incubation, 500 µL of pre-cultures were inoculated in 50-ml Erlenmeyer flasks containing 1 g of dry zeolite particles and 10 ml of MSA. The cultures were incubated at 28°C without agitation to let the cells colonize the zeolite. Every 2.5 days, 6 mL of culture medium was replaced by fresh MSA. After 3 cycles of cultivation (7.5 days), the colonized zeolite particles were collected on a sieve, rinsed twice with sterile distilled water and the residual water was absorbed by placing the sieve on paper tissues.

### Microcosm experiments

The experimental design was based on the incubation of soil microcosms either treated with atrazine or untreated, and inoculated or not with atrazine-degrading bacteria or consortia, either as free cells or biocomposite. Microcosms consisted of 1 g equivalent dry weight (dw) of soil placed in wells of sterile 24-well plates incubated at room temperature in humidity-saturated enclosures to avoid water loss. For soil equilibration, microcosms were prepared 30 days before inoculation and relative humidity was adjusted to 11%.

#### Contamination scenarios

Soil microcosms were either treated with an atrazine solution (final concentration of 1.5mg/kg dw) 15 days before inoculation (D-15), the day of inoculation (D0) or with water (NT). At D0 et D-15, the soil microcosms not receiving atrazine were complemented with ultrapure sterile water to maintain all the microcosmos at the same humidity content.

#### Mode and type of inoculation

The inoculation of soils was performed at D0 with *Pseudomonas* sp. strain ADP3 (ADP) or the consortia (CONS) either as biocomposite with zeolite as support (S) or as free living cells with no support (NS). For biocomposite inoculation, 33 mg of prepared biocomposite was mixed with the soil (equivalent to 20 g dw of zeolite per kg dw of soil). For NS condition, free cell suspensions were recovered from 33 mg biocomposite by vortexing with ultrapure sterile water (3 * 10 s). This allowed to inoculate the same cell density and composition in both S and NS conditions. Numeration of ten-fold serial dilutions of the washed biocomposite on TY agar plates indicated that the inoculum density was of 3.2 and 3.8 x10^7^ CFU per g dw of soil for the pure strain and the consortium, respectively. Controls without inoculation (NI) consisting in adding water (NS condition) or sterile zeolite (S condition) were included.

Finally the soil microcosms were adjusted to 32.6% of relative humidity with MQ water (equivalent ~60 % of the soil water holding capacity) and placed in humidity-saturated enclosures at room temperature. Soil microcosms were collected at 15 days (T1) and 45 days (T2) after inoculation for a total of 144 soil microcosms (3 inoculation types [ADP, CONS, NI] x 2 inoculation modes [S, NS] x 3 atrazine contamination scenarios [D-15, D0, NT] x 2 sampling times [T1, T2] x 4 replicates). At D-15 and D0, an additional set of 4 replicates was prepared with ^14^C-atrazine and immediately sacrificed to monitor real time herbicide mineralization in the case of atrazine contamination scenarios.

### Measurement of atrazine mineralization potential

Mineralization of atrazine was measured using microradiorespirometer following ISO14239:2017 method. A solution of atrazine containing ^14^C-ring-labelled atrazine to a final concentration of 1.5 mg and 1.4 MBq per kg dw of soil was added to soils. Whatman® 3 mm Chr paper filters soaked with a saturated solution of barite (0.37 M) were placed and sealed on the top of the 24-wells plates. During a period of incubation of 34 days, the paper filters were regularly replaced, dried and fluorography printed on Storage PhosphorScreen (Molecular Dynamics®) for 2 days along with a standard filter made of spots containing known quantities of ^14^C. The screens were finally scanned by a phosphorimager (Storm 860 Molecular Imager) and the luminescence measured due to ^14^CO_2_ trapped on the baryte paper filter during incubation was determined using the ImageQuant TL software (version 7). The luminescence was converted in radioactivity using the included standard filter and the mineralization percentage was expressed as the quantity of radioactivity measured compared to the one initially added in the microcosms. Mineralization at D-15 and D0 as well as mineralization potential at T1 and T2 of the soils microcosms were measured.

### Soil DNA extractions

Total DNA extraction was done on soil samples (250 mg dw soil; relative humidity of 32.6%) using PowerSoil DNeasy 96-well plates isolation kit (Qiagen, France) following the manufacturer’s manual. The DNA extracts were quantified using a Quant-iT PicoGreen dsDNA assay kit (Invitrogen, France) according to the manufacturer’s instructions.

### Quantification of atrazine-degrading genes

qPCR assays were conducted according to ISO17601:2016 to monitor the abundances of the atrazine-degrading genes. Prior to carry out qPCR assays, inhibition of Taq polymerase was tested by mixing soil DNA extracts with either control plasmid DNA (pGEM-T Easy Vector, Promega, France) or water. No inhibition was detected with the amount of DNA used. The qPCR assays were conducted in a total reaction volume of 15 μl including 7.5 μl master mix Takyon™ Low ROX SYBR (Eurogentec, France), 1 ng DNA extract, 1 μM of each primer, and 250 ng of T4 gene 32 (Qbiogene). Real-time PCR reactions were performed into 384-well plates using a ViiA7 (Life Technologies, USA) according to the conditions: 5 min at 95 °C, followed by 30 cycles of 15 s at 95 °C and 1 min at 60 °C (56°C for *atzD*). The calibration curves were generated using serial dilutions of plasmid DNA containing the atrazine-degrading genes (*atzA, atzB, atzC, atzD*) (Table S1) targeted by the assay. Depending on the targeted gene, PCR efficiency ranged from 91 to 117%. Each gene quantification was performed in duplicate in two independant runs.

### Assessment of soil microbial community diversity and composition

The amplicon sequencing libraries were generated with the two-step PCR (Berry et al., 2011). The first PCR reaction was performed in the final volume of 15 μl containing 7.5 μl PCR Mix, 0.375 μM from each primer, 0.05 μl T4 gene 32 (Qbiogene), and 1 ng of DNA. The hypervariable region of *16S rRNA* gene (V3–V4) was amplified by thermocycler using the primer pairs U341F and 805R (Table S1), with overhang adapters (Takahashi et al., 2014). Thermocycler conditions were 98 °C for 3 min and 25 cycles 98 °C for 30 s, 60 °C for 30 s, and 72 °C for 30 s with a final extension of 72 °C for 10 min. For the second stage, PCR products of the first step were used as a template. Besides, the PCR reaction was performed using a unique multiplex primer pair (barcode) for each sample. The reaction was performed at 40 μl volume containing 20 μl phusion master mix (Thermo Fisher Scientific), 4 μl (1µM) from the reverse barcode primer, 4 μl (1µM) from the forward barcode primer, and 6 μl from the first step PCR product. Thermocycler conditions were 98 °C for 3 min and then the eight-cycle 98 °C for 30 s, 55 °C for 30 s, and 72 °C for 30 s, with the final extension at 72 °C for 10 min. The size of PCR products of the second step was checked in agarose gel (2%) electrophoresis for visual inspection (about 630 bp). The amplicons were purified and mixed using the SequalPrep^TM^ normalization kit 96-well (Applied Biosystems™). The sequencing was performed using the MISEQ v2 kit (500 cycles).

Demultiplexing and trimming of Illumina adaptors and barcodes were done with Illumina miseq reporter software (version 2.5.1.3). Processing of raw sequences was performed using an in-house developed pipeline (https://forgemia.inra.fr/vasa/illuminametabarcoding). The partial bacterial 16S rRNA sequences were assembled using PEAR (Zhang et al., 2014) and quality-trimmed using the QIIME pipeline (Caporaso et al., 2010) and short sequences (< 400 bp) were removed. OTUs clustering was performed using VSEARCH (Rognes et al., 2016) and the sequences were classified into operational taxonomic units (OTUs) with 97% sequence identity using UCLUST (Edgar, 2010) and the SILVA database; then a representative sequence in each OTU was selected and the phylogenetic tree was constructed using FastTree (Price et al., 2010). A total of 6758 OTUs were found from the 144 samples after quality checking. Sequencing data were deposited in SRA project NCBI database (PRJNA1074952).

### Statistical analysis

Statistical analyses were conducted using R statistical software (v 4.2.3). The differences between treatments in atrazine mineralization potential and gene copy number (atrazine-degrading genes) were tested using ANOVAs followed by Tukey’s honestly significant difference test (p-value < 0.05) using the TukeyHSD() function from the *stats* package (R core team, 2023). Normality and homogeneity of the residual distributions were verified and log transformations were performed when necessary.

#### Analysis of alpha and beta-diversity

Bacterial α-diversity metrics (*i. e.* observed species, Simpson’s reciprocal and Faith’s Phylogenetic Diversity PD) and Weighted Unifrac distance (Lozupone et al., 2006) between samples were calculated based on rarefied OTU tables (12000 sequences). Analyses of the effects of the treatments and their interactions on the microbial α-diversity were performed using ANOVAs followed by multiple pairwise comparison tests using emmeans function of *emmeans* package, with Tukey correction for p-value adjustment. To detect differences in the microbial community structure between treatments, permutational multivariate analysis of variance (PermANOVA) from (Anderson, 2001) was carried out on weighted Unifrac distance matrices using the adonis function implemented in the *vegan* package (Dixon, 2003). Pairwise post hoc tests were conducted using the function pairwise.adonis from the *pairwiseAdonis* package with false discovery rate (FDR) corrections.

#### Differential abundance analysis between treatments

Low-abundance OTUs were filtered out by keeping OTUs representing more than 0.05% of the sequences across all samples and found in at least 60% of the replicates, which resulted in 412 OTUs. This filtering step allows for reducing zero counts in sequencing datasets, which can inflate the number of false positives for the differential abundance analysis. To explore which OTUs significantly changed in relative abundance between treatments, we used a generalized linear mixed model (GLMM), which allows us to infer linear regression from data that does not follow a normal distribution as abundance data typically follow a Poisson distribution and includes both treatment effects (fixed effects) and sampling effects (random effects). The analysis was performed using glmmTMB function of the *glmmTMB* package (Bolker, 2016). Multiple pairwise comparisons were performed using emmeans function (Lenth, 2019). The p-values were then adjusted using the false discovery rate (FDR) method (Benjamini and Hochberg, 1995), and only OTUs with FDR-adjusted p-values below or equal to 0.05 were considered significant.

## RESULTS

### Evaluating the efficiency and the stability of ^14^C-atrazine mineralization in soil microcosms

The assessment of atrazine mineralization dynamics was conducted in soil microcosms that were contaminated with atrazine 15 days before (D-15) or the day of inoculation (D0). Without inoculation, the soil did not exhibit any atrazine mineralization potential with less than 1.5% of atrazine mineralized after 34 days of incubation (Fig. 1A). Right after inoculation of *Pseudomonas* sp. ADP3 or of the consortia (~3.5 10^7^ CFU/ g dw of soil), mineralization started without any lag phase in both D-15 and D0 conditions. Mineralization followed a first order kinetic for all inoculated conditions, and according to the treatment between 17.2 to 54.5 % of initially added ^14^C-ring labelled atrazine was transformed into ^14^CO_2_ in 34 days. The main factors affecting the mineralization rate were the contamination scenario (D-15 or D0) and the mode of inoculation (S or NS) (ANOVA, p < 0.01) while the type of inoculation (pure strain vs consortium) had no effect. Indeed, atrazine mineralization was significantly higher in microcosms where atrazine was added at D0 (39.1 ± 2.7%) as compared to the ones where atrazine was added 15 days before (25.5 ± 1.2%) (Fig. 1B). In addition, one could observe that the microcosms amended with biocomposites presented significantly lower atrazine-mineralizing ability (26.2 ±1.7 %) than the ones inoculated with free living cells (38.4 ± 2.6%).

**Figure 1.**
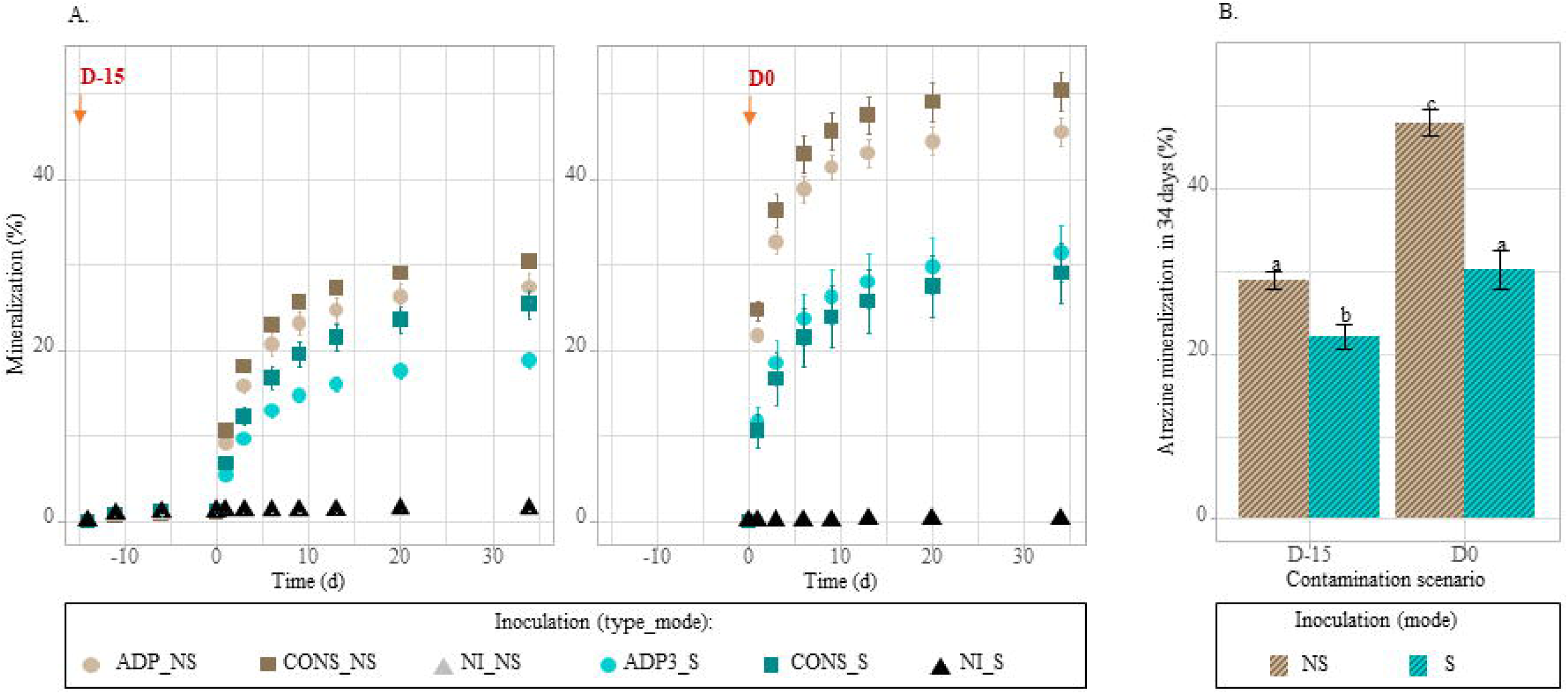
Real-time mineralisation of ^14^C-ring labelled atrazine in the soil microscoms contaminated with the herbicide 15 days before (D-15) or the day of inoculation (D0). A : Mineralisation kinetics in soil microcosms inoculated or not (NI) with the degrading strain (ADP) or consortium (CONS) as free cell (NS) or biocomposite (S). B: Cumulative percentage of atrazine mineralized after 34 days for each contamination scenario and each inoculation mode. Values are mean ± SE (n = 4) and different letters indicate statistically significant differences between conditions (ANOVA followed by a tukey’s HSD test, P < 0.05).

To assess the stability of the atrazine mineralization potential over time, the soil microcosms were analysed after 15 (T1) and 45 days (T2) of incubation (Fig. 2). First, without inoculation, no adaptation naturally appeared in the soil microcoms treated with atrazine at D-15 or D0 since atrazine mineralization potential in non-inoculated soils remained negligible (*i. e.* < 1.5% in 34 days at both T1 and T2). The mineralization potential of inoculated microcosms was maintained over the incubation period even though it signicantly decreased over time (ANOVA, F-value=322; p<0,001). Initially ranging from 17 to 54 % at D0 (Fig. 1B), mineralization capacities ranged from 10 to 22% at T1 and from 6 to 14% at T2 (Fig. 2). Interestingly, the mode of inoculation (NS or S) was the major driver of the stability of the mineralization potential (F-value=521, p<0,001). Whatever the time of sampling (T1 or T2), the contamination scenario or the type of inoculum, inoculation with biocomposites presented a mineralization potential almost twice higher than inoculation with the free living cells (18% *vs* 11% at T1, 12 *vs* 6 % at T2).

**Figure 2.**
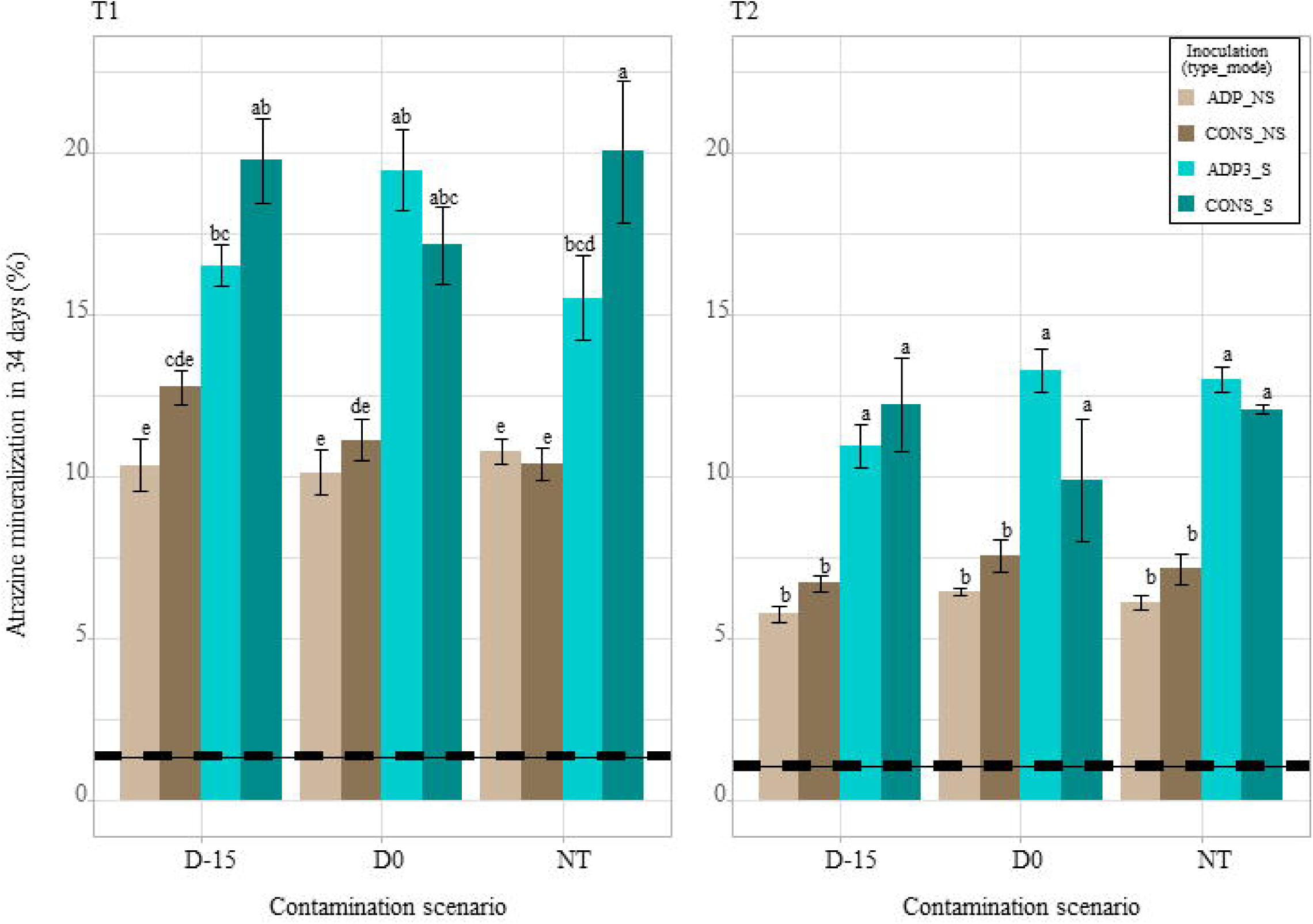
Atrazine mineralization potential of the soil microcosms measured 15 days (T1) and 34 days (T2) after inoculation with the pure strain (ADP) or the consortia (CONS) as biocomposite (S) or free-living cells (NS). Atrazine contamination scenarios are indicated on x axis: treated with atrazine 15 days before inoculation (D-15), treated the same day (D0) or not treated (NT). The mineralization potentials are expressed as cumulative percentage of ^14^CO2 emitted during 34 days compared with the amount of ^14^C-ring labelled atrazine initially supplied. Values are mean ± SE (n = 4) and different letters indicate statistically significant differences between conditions (P < 0.05). The average mineralization potential of the not-inoculated soils (NI) at T1 and T2 and its associated standard error are indicated by full and dashed lines respectively.

Finally, to a lesser extent, soils that were inoculated with the consortia tend to exhibit a higher mineralization potential compared to those inoculated with the pure strain (F-value= 9, p = 0.004). This effect is dependent on the contamination scenario, with higher mineralization potentials for inoculations with the consortium compared to inoculations with *Pseudomonas* sp. ADP3 only in D-15 contamination scenario (1.21 fold higher, p=0,003).

### Evaluating the stability of the inoculants in soil microcosms

To assess the stability of the microbial inoculants in soil, we also quantified the abundance of the sequences of atrazine-degrading genes (*atzA*, *atzC*, *atzD*). None of these *atz* genes were detected in the non-inoculated microcosms which is in accordance with the absence of mineralization potential in those microcosms. In addition, *atz* genes were quantified in all the other microcosms with sequence copy number per ng of DNA varying from 16 ± 2 to 299 ± 24 for *atzA*, from 23 ± 3 to 371 ± 20 for *atzC* and from 117 ± 7 to 1413 ± 239 for *atzD* (Fig. 3). ANOVA analyses gave the same trends for all the three *atz* genes, *i. e.* a decrease by a factor from 2.3 to 2.6 between T1 and T2. Besides the time effect, the main driver for the gene quantifications was the mode of inoculation (NS vs S) while the type of inoculum had no impact (ADP vs CONS). More precisely, the sequence numbers of *atzA*, *atzC* and *atzD* genes detected in soil inoculated with biocomposite were 3.3, 3.1 and 2.4 times higher, respectively, than in the soil inoculated with free living cells. Finally to a lesser extent, the contamination scenario affected slightly the quantity of detected *atz* genes at T1, the quantity of detected genes being lower for D0 condition, while at T2 this was only true for the soil microcosms inoculated with the consortium.

**Figure 3.**
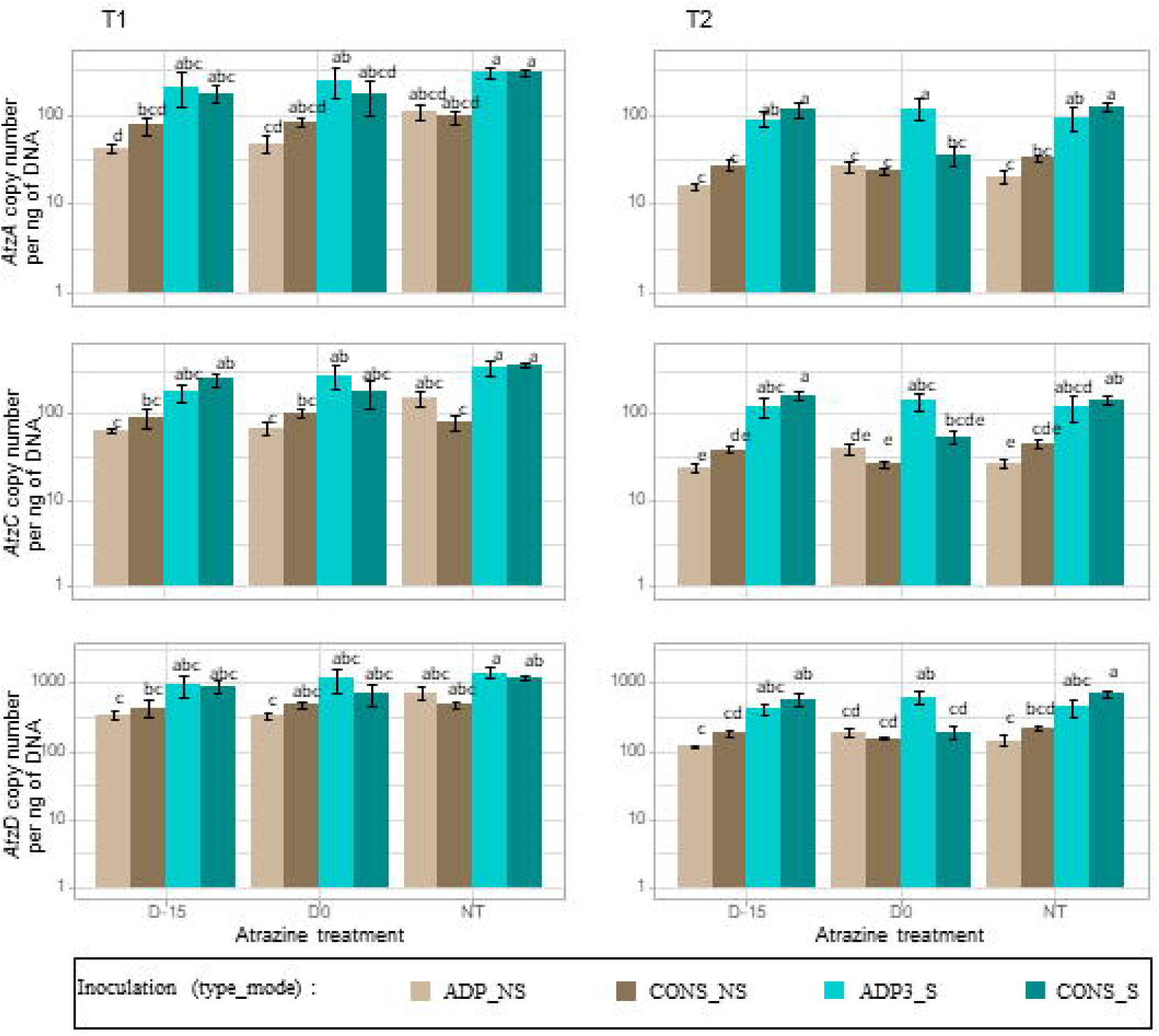
Quantification of *atzA*, *atzC* and *atzD* degrading genes in the soil microcosms at T1 and T2 by qPCR. Copy number for each gene per ng of DNA was quantified in soil microcosms inoculated with Pseudomonas sp. ADP3 (ADP) or a consortium (CONS) either inoculated as biocomoposite (S) or free-living cells (NS). The soil microcosms were subjected to different contamination scenarios: treated with atrazine 15 days before inoculation (D-15), treated the same day of inoculation (D0), or not treated (NT). Values are mean ± SE (n = 4) and different letters indicate statistically significant differences between conditions (Tukey test, p-value < 0.05).

### Evaluating the impact of treatments on the diversity of the autochtonous soil bacterial community

DNA metabarcoding was used to assess the effects of inoculation treatments with biocomposites or free-living bacteria on the native soil community at T1 and T2 sampling times. α-diversity indices relative to richness (Observed species), phylogenetic diversity (PD whole Tree) and evenness (Inverse of Simpson) were very stable whatever the treatment or the time of sampling, with coefficients of variation ranging from 1.9 % for richness to 6 % for the evenness indicating a quantitatively small impact of inoculation treatments on bacterial diversity. Despite this stability, ANOVA analysis revealed significant differences due to time of sampling and treatments (p<0.05) (Fig. S1). The sampling time was the major driver of all α-diversity indices. Significant impact of the treatments on α-diversity indices were observed only for T2 sampling time (Table S2). Indeed, soils supplemented with zeolite presented higher richness (2240 ± 6 *vs* 2207 ± 7 observed species) and phylogenetic diversity (231 ±1 *vs* 228 ± 1 AU) and thoses effects were even greater when the consortium was inoculated. Concerning evenness, it was impacted by the atrazine contamination scenario showing a higher evenness when atrazine was added at D-15 as compared to the uncontaminated soils (356 ± 5 *vs* 341 ± 4 AU), this effect being higher in presence of zeolite (359 ± 7 *vs* 334 ± 5 AU).

### Evaluating the impact of the treatments on the structure of autochtonous soil bacterial community

We then investigated the impact of the treatments on the microbial community structure using Principal Coordinates Analysis (PCoA) based on weighted Unifrac distances analysis (Fig 4). Our results showed that at both time of sampling (T1 and T2), the structure of the microbial community was significantly impacted by the different inoculation treatments (triple interaction between contamination scenarios, inoculation type and inoculation mode) with the strongest effect, attributed at both timepoints to the *Type-by-Contamination Scenario* interaction (Table S3). To asess the size of these effects and identify OTUs affected by the different inoculation treatments, we performed a differential abundance analysis of the microbial community composition at each sampling time and under different atrazine contamination scenarios (*i. e*. D-15, D0, and NT) (Fig. 5 and Fig. S2; Table S4).. Our analysis showed a higher number of affected OTUs at T2 sampling time compared to T1 (between 8% and 55% at T2 and between 9% and 19% at T1 depending on the atrazine contamination scenarios). By the end of the experiment, we found that the native microbial community was differentially affected depending on the atrazine contamination scenario, with a higher number of impacted OTUs when soil microcosms were inoculated concomitantly to the atrazine contamination (D0) (Fig. 5). As such, pairwise comparisons between treatments showed that in total 18%, 55%, and 8% of the dominant OTUs were significantly affected by the type of inoculum (single species *vs* consortium) and the inoculation mode (zeolite *vs* free-living) in D-15, D0, and NT communities, respectively (Fig. 5; FDR-adjusted *P ≤ 0.05*). To differentiate the relative importance of the effect of the type of inoculum and the inoculation mode, we further focused on their impact on the relative abundances of the bacterial community in the D0 contamination scenario. Thus, we observed a more pronounced impact of the bacterial consortium on the community composition (CONS vs NI = 26% affected OTUs) compared to the single strain (ADP vs NI = 1.21% affected OTUs) during the D0 contamination scenario. These observations were supported by a significant disparity in the community composition between the consortium and single strain (CONS vs ADP3 = 29.8% affected OTUs). However, these differences were not observed at D-15 and NT contamination scenarios, indicating that the effect of the consortium on the native community was mediated by atrazine contamination. The differential abundances analysis revealed that members of Actinobacteria significantly decreased in soil microcosms inoculated with the bacterial consortium, while Bacteroides and Acidobacteria exhibited an increase in their relative abundances in D0 communities (*i.e.* CONS > ADP and CONS > NI; Fig. 6). When comparing ADP-S and ADP-NS treatments in D0 communities, we found that 17.7% of the most abundant OTUs were significantly impacted. Interestingly, there were also significant shifts in the relative abundances of 14% of the OTUs within the D0 community observed between NI-S and NI-NS, while no differences were detected between these treatments in the NT community, indicating an interplay effect between atrazine contamination scenario and zeolite on the composition of the microbial community. For example, we found that the use of zeolite significantly increased the relative abundances of OTUs belonging to Bacteroidetes and Gammaproteobacteria (*i. e.* ADP < NI-NS and ADP-NS < ADP-S). In contrast, OTUs from Actinobacteria group were negatively impacted by the zeolite treatment (Fig.6).

**Figure 4.**
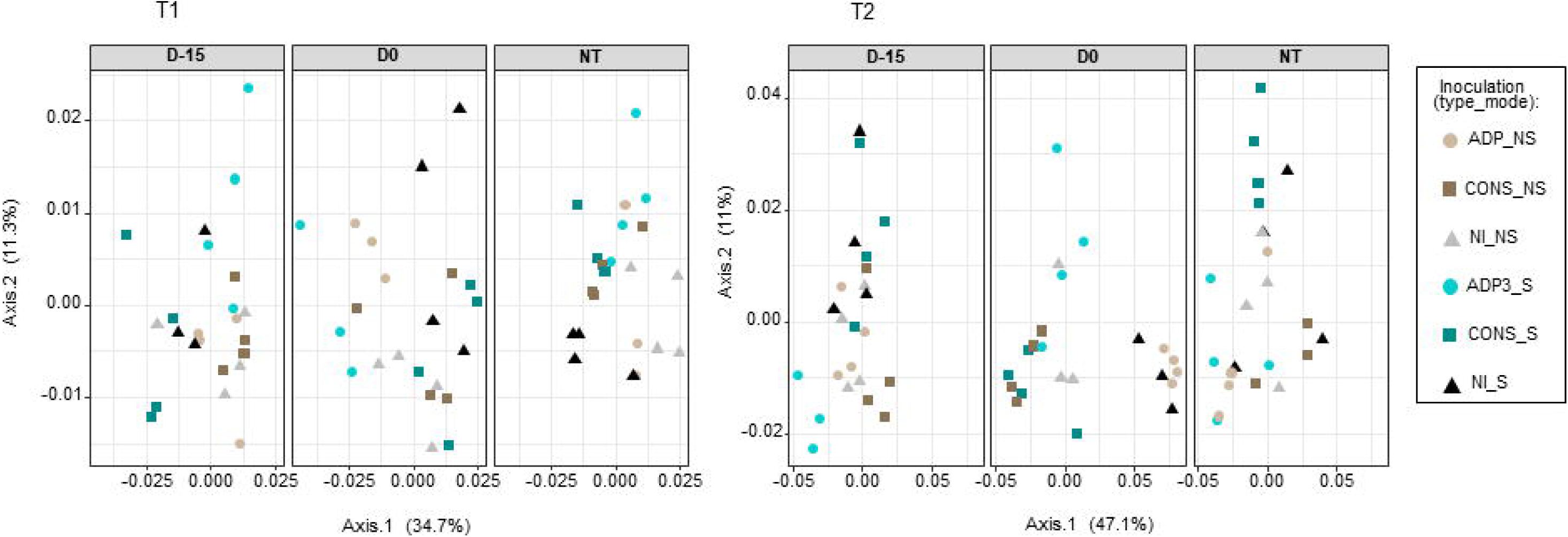
Bacterial community structure of the soil microcosms inoculated with *Pseudomonas* sp. ADP3 (ADP) or a consortium (CONS) either inoculated as biocomposite (S) or free-living cells (NS) after 15 (T1) or 34 (T2) days of inoculation. Dissimilarity in community structure was measured with the weighted UniFrac distance. The soil microcosms were subjected to different atrazine contamination scenarios: treated with atrazine 15 days before inoculation (D-15), treated the same day of inoculation (D0), or not treated (NT). The percentage of the total variation was showed in brackets on each axis.

**Figure 5.**
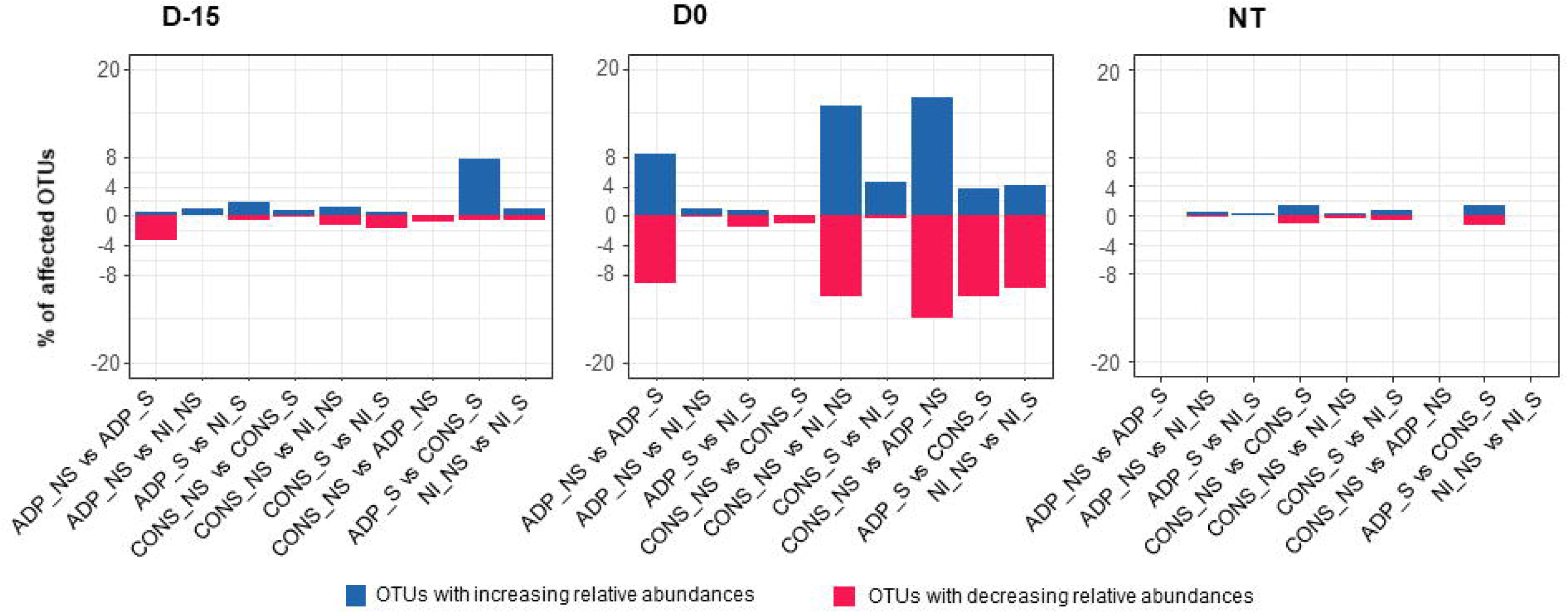
Changes in the relative abundances of the most abundant OTUs between soil microcosms inoculated with *Pseudomonas* sp. ADP3 (ADP) or a consortium (CONS) as biocomposites or free-living cells. Differentially abundant OTUs were identified using a generalized linear mixed model. The soil microcosms were exposed to different atrazine contamination scenarios: treated with atrazine 15 days before inoculation (D-15), treated the same day of inoculation (D0), or not treated (NT). The percentage of OTUs exhibiting significantly increasing/decreasing relative abundances for each pairwise (where vs means > or <) are presented in blue and red, respectively.

**Figure 6.**
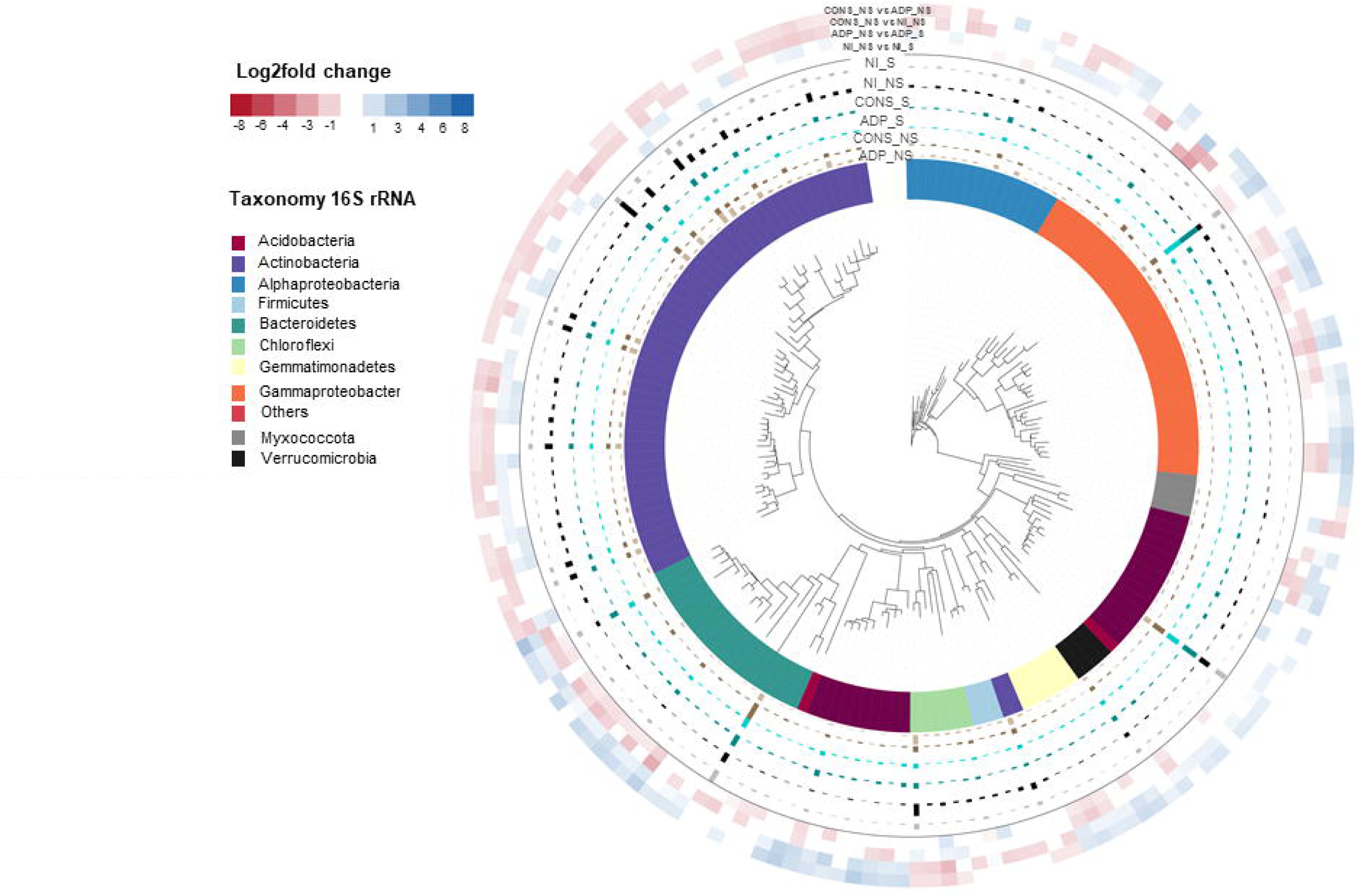
Taxonomic assignment and distribution of significantly differentially abundant OTUs between treatments in the D0 contamination scenario at T2 sampling time. Outer rings show a heatmap of OTUs with significantly increasing/decreasing relative abundances between CONS and ADP, CONS and NI, ADP and ADP-S, and NI and NI-NS treatments (where vs means > or <). Changes in the relative abundances, expressed as the Log2fold change, are represented by the blue-to-red color. Bar plots represents the mean relative abundances of OTUs in each treatment. The affiliation of OTUs at the phylum or class levels is indicated by different colors on the internal ring.

## DISCUSSION

Here, we show that the inoculation of atrazine degrading bacteria in soil, either as a single strain or as a consortium, enables the immediate degradation of atrazine after its application. Indeed, free-living inoculated bacteria mineralized up to 50% of the molecule in less than 30 days. Other studies have already shown that albeit with varying degrees of success (Topp, 2001; Rousseaux et al., 2003; Chelinho et al., 2012). Our work also underlines that a classic bioaugmentation strategy, *i. e.* using a pure strain in liquid culture without the addition of atrazine, is not optimal and that the mineralization function is difficult to stabilize. The mineralisation potential 45 days after inoculation decreased to less than 8% probably related to the difficulty of the inoculum to establish in soil, as indicated by quantification of the inoculum by qPCR. Therefore, although mineralization remains quantifiable after 45 days, its reduced efficiency prevents an optimal removal of further atrazine influxes before the molecule is transferred to the other compartments. These results are in line with the literature, which underlines the difficulty of implanting a new function in a soil through the establishment of a microbial inoculum which must adapt to both abiotic constraints (reviewed by Cycon et al., 2017) and biotic competition (potent as an invader)(Mallon et al., 2015).

To improve the strains’ ability to establish and express their mineralization potential, we tested whether the addition of atrazine at the time of inoculation would be beneficial. Various studies have shown that providing the inoculated strains with a substrate that confers them a temporarily available niche in soil can decrease the immediate competition with the indigenous microbiota (Causevic et al., 2024). In our case, the addition of atrazine had surprisingly little effect on the maintenance of the strain and its function over time, after 15 and 45 days. Piutti et al. (2002) previously showed that the increase of the inoculum density related to the substrate addition was only transient and concomitant with the exponential phase of the substrate mineralization (Piutti et al., 2002). In our case, the exponential phase of mineralization occurs in the first 5 days after inoculation, and although this probably led to an increase in the inoculum density, it did not result into a better maintenance after 15 and 45 days.

We also tested the hypothesis that inoculating a consortium rather than a single strain would result in greater stability of the inoculum and hence the function. Some studies suggested that complex communities, through higher levels of genetic complexity, metabolism, cellular communication and cooperation, are more likely to settle down and are more efficient for biodegradation (Pileggi et al., 2020). This hypothesis was not verified in our case, eventhough a slight increase of mineralization was observed with the consortia as compared to the single strain.

Confirming our second hypothesis, we showed that the most effective strategy for improving bioaugmentation stability and efficiency was to inoculate not as free-living cells but as a biocomposite, using zeolite as the carrier material. Inoculation of the consortium or *Pseudomonas* sp. ADP3 grown as biofilms on zeolite particles led to better survival of the inocula compared to free-living cells at both short and long term (15 and 45 days after inoculations) and, in turn, to a greater stability of atrazine mineralization potential over time (Fig. 2 and 3). The advantage of this approach over others, such as substrate addition or repeated inoculations (Newcombe and Crowley, 1999), is that the carrier material not only provides a niche for the inocula that are not directly in competition with soil microorganisms, but also constitutes a protective environment that buffers the physico-chemical environment surrounding the bacteria from the soil environment (Gorodylova et al., 2021). Another potential advantage of zeolite over other carrier materials is itshigh cation exchange capacity (CEC), which may enhance its ability to retain pollutants such as pesticides. In addition to improving the survival of the degraders, it could concentrate the pollutants to be remedied. Recent studies have tested bioremediation strategies of atrazine with different biocomposite formulations such as activated carbon (Zhang et al., 2024), alginate beads (Pan et al., 2023) or encapsulation (Gal et al., 2021), and they all displayed significant improvements of inoculation of biocomposites compared to free-living cells. One limitation, previouslydemonstrated in the case of alginate, is that using a carrier material can limit access to the substrate by limiting the dispersion of the inoculum. This phenomenon seems to happen in our case because, right after inoculation at D0, the immobilized cells showed a lower mineralization potential than the free-living cells for which the inoculation led to the simultaneous dispersion of both the inocula and the applied atrazine throughout the entire soil volume. This highlights the importance of the colocation of the pollutant and the microbial degraders for an effective bioremediation (Pinheiro et al., 2015).

Another important point to evaluate when proposing a bioremediation strategy based on the use of biocomposites is to monitor the absence of long-term unintended ecotoxicological effects on soil ecosystems. Here, we assessed the impact of both the inoculation type (single strain versus consortium) and the inoculation mode (zeolithe as a support versus free living cells) on the authochtonous soil bacterial community diversity and structure, under three atrazine contamination scenarios. We observed a slight impact of the use of biocomposites on the diversity of the soil bacterial community with an increase in bacterial phylogenetic diversity and richness when using the consortium-based biocomposites at T2. The main impact on soil microbial community composition was also observed at T2 and related to an interplay effect between atrazine contamination at D0 and the inoculation treatments, with a relatively strong impact of the consortium compared to the single strain. Globally, inoculating the consortium-based biocomposites when atrazine was added simultaneously induced a shift in the structure of the soil bacterial community, possibly because adding atrazine improved the competitiveness of the consortium, and hence its impact. We however did not evaluate the survival at longer term of the atrazine degraders and the potential functional impacts on soil ecosystem services that were caused by the soil bacterial community modifications.

## Conclusion

Altogether, we demonstrated here the increased stability and efficiency over time of zeolite-based biocomposites for the bioremediation of atrazine contamination in an agricultural soil. We also showed that the overall impact of this technique on the soil autochthonous bacterial community structure and diversity is relatively mild if the contamination is not concomitant with the inoculation. Further researches are needed to evaluate in more depth the ecotoxicological impacts of this bioaugmentation technique on soil ecosystem services to validate its use as a safe and eco-friendly technique for the cost-effective and sustainable detoxification of pesticide contaminants in soils.

## AUTHOR CONTRIBUTIONS

Conceptualization: MD-L, FM-L, AS; Formal Analysis: SA, MD-L, SR, NR; Funding Acquisition: CM, FM-L, AS; Investigation: SA, MD-L, SR, NR, AS; Methodology: MD-L, SR, NR; Supervision; MD-L, SR, AS; Validation: AS; Visualization: MD-L, SR; Writing – original draft: SA, AS; Writing – review and editing: MD-L, FM-L, CM, SR

## FUNDING SOURCES

This work was supported by the French ANR [ANR-21-CE04-0018 EPURSOL, 2021].

## Notes

### Competing Interest Statement

The authors have declared no competing interest.

## REFERENCES

Anderson, MJ. 2001. A new method for non-parametric multivariate analysis of variance. Austral. Ecol. 46:26–32.

Benjamini, Y., & Hochberg, Y. 1995. Controlling the false discovery rate: a practical and powerful approach to multiple testing. J. R. STAT. SOC. B. 57: 289–300.

Berry, D., Ben, M. K., Wagner, M. & Loy, A. 2011. Barcoded primers used in multiplex amplicon pyrosequencing bias amplification. Appl. Environ. Microbiol. 77: 7846–7849.

Billet, L., Devers, M., Rouard, N., Martin-Laurent, F. & Spor, A. 2019. Labour sharing promotes coexistence in atrazine-degrading bacterial communities. Sci. Rep. 9(1): 18363.

Billet, L., Devers-Lamrani, M., Serre, R. F., Julia, E., Vandecasteele, C., Rouard, N., & Spor, A. 2021. Complete genome sequences of four atrazine-degrading bacterial strains, Pseudomonas sp. strain ADPe, Arthrobacter sp. strain TES, Variovorax sp. strain 38R, and Chelatobacter sp. strain SR38. Microbiol. Resour. Announc. 10 (1): e01080–20.

Bolker, B. 2016. Getting started with the glmmTMB package. Vienna, Austria: R Foundation for Statistical Computing. software. https://cran.r-project.org/web/packages/glmmTMB/vignettes/glmmTMB.

Borges, I. L., Forsyth, L. Z., Start, D., & Gilbert, B. 2019. Abiotic heterogeneity underlies trait-based competition and assembly. J. Ecol. 107(2): 747–756.

Caporaso, JG., Kuczynski, J., Stombaugh, J., Bittinger, K., Bushman, FD., Costello. EK, et al. 2010. QIIME allows analysis of high-throughput community sequencing data. Nat. Methods. 7:335–6.

Carles, L., Martin-Laurent, F., Devers, M., Spor, A., Rouard, N., Beguet, J. & Batisson, I. 2021. Potential of preventive bioremediation to reduce environmental contamination by pesticides in an agricultural context: a case study with the herbicide 2, 4-D. J. Hazard. Mater. 416, 125740.

Causevic, S., Dubey, M., Morales, M., Salazar, G., Sentchilo, V., Carraro, N., & van der Meer, J. R. 2024. Niche availability and competitive loss by facilitation control proliferation of bacterial strains intended for soil microbiome interventions. Nat. Commun. 15(1), 2557

Chen Y., Jiang Z., Wu D., Wang H., Li J., Bi M., & Zhang Y. 2019. Development of a novel bio-organic fertilizer for the removal of atrazine in soil. J. Environ. Manage. 233: 553–560.

Chelinho, S., Moreira-Santos, M., Silva, C., Costa, C., Viana, P., Viegas, C. A. & Sousa, J. P. 2012. Semifield testing of a bioremediation tool for atrazine-contaminated soils: Evaluating the efficacy on soil and aquatic compartments. Environ.Toxicol.Chem. 31(7), 1564–1572.

Cycon, M., Mrozik, A., & Piotrowska-Seget, Z. 2017. Bioaugmentation as a strategy for the remediation of pesticide-polluted soil: A review. Chemosphere, 172, 52–71.

De Souza, M. L., Sadowsky, M. J., & Wackett, L. P. 1996. Atrazine chlorohydrolase from Pseudomonas sp. strain ADP: gene sequence, enzyme purification, and protein characterization. J. Bacteriol. 178(16), 4894–4900.

Dixon, P. 2003. VEGAN, a package of R functions for community ecology. J. veg. sci. 14(6): 927–930.

Dominguez-Garay, A., Boltes, K., & Esteve-Nunez, A. 2016. Cleaning-up atrazine-polluted soil by using microbial electroremediating cells. Chemosphere. 161: 365–371.

Edgar, R. C. 2010. Search and clustering orders of magnitude faster than BLAST. Bioinform. 26(19): 2460–2461.

Gal, R., Perez-Lapid, N., Zvulunov, Y., & Radian, A. 2021. Layer-by-Layer Encapsulation of Herbicide-Degrading Bacteria for Improved Surface Properties and Compatibility in Soils. Polymers. 13(21), 3814.

Gorodylova, N., Michel, C., Seron, A., Joulian, C., Delorme, F., Bresch, S., & Michel, K. 2021. Modified zeolite-supported biofilm in service of pesticide biodegradation. Environ. Sci. Pollut. Res. 28: 45296–45316.

ISO 17601:2016. Soil quality — Estimation of abundance of selected microbial gene sequences by quantitative PCR from DNA directly extracted from soil. https://www.iso.org/standard/60106.html

ISO 14239:2017. Soil quality - Laboratory incubation systems for measuring the mineralization of organic chemicals in soil under aerobic conditions.

Jia, W., Li, N., Yang, T., Dai, W., Jiang, J., Chen, K. & Xu, X. 2021. Bioaugmentation of atrazine-contaminated soil with Paenarthrobacter sp. strain AT-5 and its effect on the soil microbiome. Front. microbiol. 3763.

Jiang, D., Huang, Q., Cai, P., Rong, X. & Chen, W. 2007. Adsorption of Pseudomonas putida on clay minerals and iron oxide. ColloidsSurf B: Biointerfaces. 54:217–221.

Kuldeyev, E., Seitzhanova, M., Tanirbergenova, S., Tazhu, K., Doszhanov, E., Mansurov, Z. & Berndtsson, R. 2023. Modifying Natural Zeolites to Improve Heavy Metal Adsorption. Water. 15(12): 2215.

Lenth, R., Singmann, H., Love, J., Buerkner, P., & Herve, M. 2019. Emmeans: estimated marginal means, aka least-squares means (Version 1.3. 4). Emmeans Estim. Marg. Means Aka Least-Sq. Means https://CRAN.R-project.org/package=emmeans.

Lozupone, C., Hamady, M., & Knight, R. 2006. UniFrac–an online tool for comparing microbial community diversity in a phylogenetic context. BMC bioinformatics. 7(1): 1–14.

Mallon, C. A., Van Elsas, J. D., & Salles, J. F. 2015. Microbial invasions: the process, patterns, and mechanisms. Trends. Microbiol. 23(11), 719–729.

Mandelbaum, R. T., Wackett, L. P., & Allan, D. L. 1993. Mineralization of the s-triazine ring of atrazine by stable bacterial mixed cultures. Appl. Environ. Microbiol. 59(6), 1695–1701.

Mohsin, M.Z., Huang, J., Hussain, M.H., Zaman, W.Q., Liu, Z., Rehman, S., Zhuang, Y., Guo, M. and Mohsin, A. 2023. Revolutionizing bioremediation: Advances in zeolite-based nanocomposites. Coord. Chem. Rev. 491: 215253.

Newcombe, D. A., & Crowley, D. E. 1999. Bioremediation of atrazine-contaminated soil by repeated applications of atrazine-degrading bacteria. Appl. Microbiol. Biotechnol. 51, 877–882.

Jia, W., Li, N., Yang, T., Dai, W., Jiang, J., Chen, K. & Xu, X. 2021. Bioaugmentation of atrazine-contaminated soil with Paenarthrobacter sp. strain AT-5 and its effect on the soil microbiome. Front. microbiol : 3763.

Jiang, D., Huang, Q., Cai, P., Rong, X. & Chen, W. 2007. Adsorption of Pseudomonas putida on clay minerals and iron oxide. ColloidsSurf B: Biointerfaces, 54:217–221.

Pan, Z., Wu, Y., Zhai, Q., Tang, Y., Liu, X., Xu, X., & Zhang, H. 2023. Immobilization of bacterial mixture of Klebsiella variicola FH-1 and Arthrobacter sp. NJ-1 enhances the bioremediation of atrazine-polluted soil environments. Front.Microbiol. 14, 1056264.

Pileggi, M., Pileggi, S. A., & Sadowsky, M. J. 2020. Herbicide bioremediation: from strains to bacterial communities. Heliyon, 6(12).

Pinheiro, M., Garnier, P., Beguet, J., Laurent, F. M., & Gonod, L. V. 2015. The millimetre-scale distribution of 2, 4-D and its degraders drives the fate of 2, 4-D at the soil core scale. Soil Biol. Biochem. 88, 90–100.

Piutti, S., Hallet, S., Rousseaux, S., Philippot, L., Soulas, G., & Martin-Laurent, F. 2002. Accelerated mineralisation of atrazine in maize rhizosphere soil. Biol. Fertil. Soils. 36, 434–441.

Rognes, T., Flouri, T., Nichols, B., Quince, C. and Mahe, F. 2016. VSEARCH: a versatile open source tool for metagenomics. PeerJ. 4: e2584.

Rostami, S., Jafari, S., Moeini, Z., Jaskulak, M., Keshtgar, L., Badeenezhad, A. & Dehghani, M. 2021. Current methods and technologies for degradation of atrazine in contaminated soil and water: A review. Environ. Technol. Innov. 24: 102019.

Rousseaux, S., Hartmann, A., Lagacherie, B., Piutti, S., Andreux, F., & Soulas, G. 2003. Inoculation of an atrazine-degrading strain, Chelatobacter heintzii Cit1, in four different soils: effects of different inoculum densities. Chemosphere. 51(7), 569–576.

Takahashi, S., Junko, T., Kaori, N., Takayoshi, H. & Miyuki, N. 2014. Development of a prokaryotic universal primer for simultaneous analysis of Bacteria and Archaea using next-generation sequencing. PLoS ONE. 9: e105592.

Topp, E. 2001. A comparison of three atrazine-degrading bacteria for soil bioremediation. Biol. Fertil. Soils. 33, 529–534.

Sanchez, V., Lopez-Bellido, J., Rodrigo, M. A. & Rodríguez, L. 2019. Enhancing the removal of atrazine from soils by electrokinetic-assisted phytoremediation using ryegrass (Lolium perenne L.). Chemosphere. 232: 204–212.

Satriani, A., Belviso, C., Lovelli, S., di Prima, S., Coppola, A., Hassan, S. B., & Comegna, A. 2024. Impact of a synthetic zeolite mixed with soils of different pedological characteristics on soil physical quality indices. Geoderma. 451, 117084

Struthers, J. K., Jayachandran, K., & Moorman, T. 1998. Biodegradation of atrazine by Agrobacterium radiobacter J14a and use of this strain in bioremediation of contaminated soil. Appl. Environ. Microbiol. 64(9): 3368–3375

Szatanik-Kloc, A., Szerement, J., Adamczuk, A., & Jozefaciuk, G. 2021. Effect of low zeolite doses on plants and soil physicochemical properties. Materials, 14(10), 2617.

Zhang, J., Kobert, K., Flouri, T. and Stamatakis, A. 2014. PEAR: a fast and accurate Illumina Paired-End reAd mergeR. Bioinformatics. 30:614–20.

Zhang, T., & Zhang, H. 2022. Microbial consortia are needed to degrade soil pollutants. Microorganisms. 10(2): 261.

Zhang, B., Zhang, J., Wang, Y., Qu, J., Jiang, Z., Zhang, X. & Zhang, Y. 2024. Biodegradation of atrazine with biochar-mediated functional bacterial biofilm: Construction, characterization and mechanisms. J.Hazard.Mate. 465, 133237.

Zhao, X., Wang, L., Ma, F. & Yang, J. 2018. Characterisation of an efficient atrazine-degrading bacterium, Arthrobacter sp. ZXY-2: an attempt to lay the foundation for potential bioaugmentation applications. Biotechnol. biofuels. 11(1): 1–10.

Zheng, X., Arps, PJ. & Smith, RW. 2001. Adhesion of two bacteria onto dolomite and apatite: their effect on dolomite depression in anionic flotation. Int. J. Miner. Process. 62:159–172.

Zhu, J., Fu, L., Jin, C., Meng, Z., & Yang, N. 2019. Study on the isolation of two atrazine-degrading bacteria and the development of a microbial agent. Microorganisms. 7(3): 80.

