## Supplementary Files for "Assessing the efficiency and the side effects of atrazine-degrading biocomposites amended to atrazine-contaminated soil"

**Supplementary Tables**

**Table S1. Sequences of the primer pairs used in this study**

| **Target gene** | **Annealing (℃)** | **sequences** |
| --- | --- | --- |
| Sp6-T7 | 60 | 5ʹATTTAGGTGACACTATAG3ʹ 5ʹTAATACGACTCACTATAGGG3ʹ |
| *atzA* | 60 | 5ʹACACCCACCTCACCATAGACC3ʹ 5ʹCGGGCGTCAATTCTATGAC3ʹ |
| *atzB* | 60 | 5ʹAGGGTGTTAGGTGGTGAAC3ʹ 5ʹCACCACTGTGCTGTGGTAGA3ʹ |
| *atzC* | 60 | 5ʹGCTCACATGCAGGTACTCCA3ʹ 5ʹTCCCCCAACTAAATCACAGC3ʹ |
| *atzD* | 56 | 5ʹTCCCACCTGACATCACAAAC3ʹ 5ʹGGGTCTCGAGGTTTGATTG3ʹ |
| U341F-805R | 60 | 5′CCTACGGGRSGCAGCAG3′ 5′GACTACCAGGGTATCTAAT3′ |

**Table S2.** **Effects of the experimental factors and their interaction on bacterial alpha diversity at each sampling time**. Impact of inoculation mode (Mode) and type (Type) under three atrazine contamination scenarios (Cont. Scen.) on phylogenetic diversity (A), observed species (B), and inverse Simpson index (C).

**A**

| **Sampling time T1** |  | |  | |  |  |  |
| --- | --- | --- | --- | --- | --- | --- | --- |
| Treatment | Df | | Sum sq | | Mean sq | F | P value |
| Type | | 2 | | 44.5 | 22.26 | 1.22 | 0.30 |
| Contamination Scenario | | 2 | | 15.7 | 8.77 | 0.43 | 0.652 |
| Mode | | 1 | | 0.6 | 0.610 | 0.033 | 0.856 |
| Type × Cont. Scen. | | 4 | | 115.8 | 28.95 | 1.58 | 0.192 |
| Type × Mode | | 2 | | 1.2 | 0.609 | 0.033 | 0.967 |
| Cont. Scen. × Mode | | 2 | | 3.2 | 1.619 | 0.089 | 0.915 |
| Type × Cont. Scen. × Mode | | 4 | | 84.6 | 21.149 | 1.161 | 0.34 |
| Residuals | | 49 | | 892.6 | 18.217 |  |  |
| **Sampling time T2** | |  | |  |  |  |  |
| Type | | 2 | | 96.1 | 48.05 | 2.391 | 0.010 |
| Contamination Scenario | | 2 | | 18.4 | 9.21 | 0.458 | 0.635 |
| Mode | | 1 | | 235.4 | 235.42 | 11.71 | 0.0012** |
| Type × Cont. Scen. | | 4 | | 24.6 | 6.16 | 0.30 | 0.872 |
| Type × Mode | | 2 | | 202.1 | 101.04 | 5.02 | 0.010* |
| Cont. Scen. × Mode | | 2 | | 13.6 | 6.81 | 0.33 | 0.714 |
| Type × Cont. Scen. × Mode | | 4 | | 216.7 | 54.17 | 2.69 | 0.041* |
| Residuals | | 50 | | 1004.8 | 20.10 |  |  |

**B**

| **Sampling time T1** |  | |  | |  |  |  |
| --- | --- | --- | --- | --- | --- | --- | --- |
| Treatment | Df | | Sum sq | | Mean sq | F | P value |
| Type | | 2 | | 6650 | 3325 | 2.26 | 0.11 |
| Contamination Scenario | | 2 | | 2176 | 1088 | 0.742 | 0.481 |
| Mode | | 1 | | 572 | 572 | 0.39 | 0.53 |
| Type × Cont. Scen. | | 4 | | 8080 | 2020 | 1.37 | 0.25 |
| Type × Mode | | 2 | | 1949 | 974 | 0.66 | 0.519 |
| Cont. Scen. × Mode | | 2 | | 606 | 303 | 0.20 | 0.814 |
| Type × Cont. Scen. × Mode | | 4 | | 1293 | 3233 | 2.20 | 0.08 |
| Residuals | | 49 | | 71904 | 1467 |  |  |
| **Sampling time T2** | |  | |  |  |  |  |
| Type | | 2 | | 1015 | 5079 | 3.13 | 0.04* |
| Contamination Scenario | | 2 | | 1064 | 532 | 0.34 | 0.70 |
| Mode | | 1 | | 17245 | 17245 | 11.24 | 0.0015** |
| Type × Cont. Scen. | | 4 | | 1578 | 394 | 0.25 | 0.90 |
| Type × Mode | | 2 | | 1289 | 6446 | 4.20 | 0.02* |
| Cont. Scen. × Mode | | 2 | | 296 | 148 | 0.096 | 0.90 |
| Type × Cont. Scen. × Mode | | 4 | | 7097 | 1774 | 1.15 | 0.34 |
| Residuals | | 50 | |  |  |  |  |

**C**

| **Sampling time T1** |  | |  | |  |  |  |
| --- | --- | --- | --- | --- | --- | --- | --- |
| Treatment | Df | | Sum sq | | Mean sq | F | P value |
| Type | | 2 | | 1079 | 539.5 | 1.38 | 0.25 |
| Contamination Scenario | | 2 | | 2236 | 118 | 2.87 | 0.06 |
| Mode | | 1 | | 94 | 93.5 | 0.24 | 0.62 |
| Type × Cont. Scen. | | 4 | | 987 | 246.9 | 0.63 | 0.63 |
| Type × Mode | | 2 | | 1714 | 856.9 | 2.20 | 0.12 |
| Cont. Scen. × Mode | | 2 | | 1513 | 756.4 | 1.94 | 0.15 |
| Type × Cont. Scen. × Mode | | 4 | | 2134 | 533.5 | 1.37 | 0.25 |
| Residuals | | 49 | | 19037 | 388.5 |  |  |
| **Sampling time T2** | |  | |  |  |  |  |
| Type | | 2 | | 91 | 45.3 | 0.137 | 0.87 |
| Contamination Scenario | | 2 | | 2268 | 1134 | 3.42 | 0.04* |
| Mode | | 1 | | 3 | 3.4 | 0.010 | 0.91 |
| Type × Cont. Scen. | | 4 | | 3960 | 989.9 | 2.98 | 0.02* |
| Type × Mode | | 2 | | 2479 | 1239.7 | 3.74 | 0.03* |
| Cont. Scen. × Mode | | 2 | | 2877 | 1438 | 4.34 | 0.018* |
| Type × Cont. Scen. × Mode | | 4 | | 3759 | 939.5 | 2.83 | 0.03* |
| Residuals | | 50 | | 1656 | 331.3 |  |  |

Significant codes: 0 ‘***’ 0.001 ‘**’ 0.01 ‘*’ 0.05 ‘.’ 0.1 ‘ ’ 1

| **Table S3. Analysis of differences between treatments based on the weighted Unifrac distances for each sampling time.** Multivariate permutational analyses of variance (PERMANOVA) results assessing the effects of inoculation mode (Mode) and type (Type) under three atrazine contamination scenarios (Cont. Scen.) on in the bacterial community structure. |  |  |  |  | **SAMPLING TIME = T2** |
| --- | --- | --- | --- | --- | --- |
|  | | |  |  | Permutation test for adonis under reduced model |
| **Sampling time T1** \| \| \| \| \| \| \|; Treatments \| Df \| \| Sum sq \| \| R^2^ \| F \| P value; Type \| \| 2 \| \| 0.002790 \| 0.04995 \| 2.3893 \| 0.012 ^*^; Mode \| \| 1 \| \| 0.000596 \| 0.01067 \| 1.0205 \| 0.357; Contamination Scenario \| \| 2 \| \| 0.001950 \| 0.03490 \| 1.6697 \| 0.072; Type × Mode \| \| 2 \| \| 0.003006 \| 0.003006 \| 2.5742 \| 0.006 ^**^; Type × Cont. Scen. \| \| 4 \| \| 0.009133 \| 0.16348 \| 3.9104 \| 0.001 ^***^; Mode × Cont. Scen. \| \| 2 \| \| 0.002688 \| 0.04811 \| 2.3015 \| 0.007 ^**^; Type × Mode × Cont. Scen. \| \| 4 \| \| 0.007091 \| 0.12693 \| 3.0360 \| 0.001 ^***^; Residual \| \| 49 \| \| 0.028611 \| 0.51215 \| \|; Total \| \| 66 \| \| 0.055865 \| 1.00000 \| \|; **Sampling time T2** \| \| \| \| \| \| \|; Type \| \| 2 \| \| 0.007026 \| 0.04127 \| 2.6678 \| 0.012 ^*^; Mode \| \| 1 \| \| 0.012162 \| 0.07144 \| 9.2358 \| 0.001 ^***^; Contamination Scenario \| \| 2 \| \| 0.013563 \| 0.07967 \| 5.1499 \| 0.001 ^***^; Type × Mode \| \| 2 \| \| 0.008941 \| 0.05252 \| 3.3947 \| 0.003 ^**^; Type × Cont. Scen. \| \| 4 \| \| 0.037295 \| 0.21908 \| 7.0804 \| 0.001 ^***^; Mode × Cont. Scen. \| \| 2 \| \| 0.009254 \| 0.05436 \| 3.5136 \| 0.002 ^**^; Type × Mode × Cont. Scen. \| \| 4 \| \| 0.016155 \| 0.09490 \| 3.0670 \| 0.002 ^**^; Residual \| \| 50 \| \| 0.065842 \| 0.38677 \| \|; Total \| \| 67 \| \| 0.170237 \| 1.00000 \| \| | |  |  |  |  |

Significant codes : 0 ‘***’ 0.001 ‘**’ 0.01 ‘*’ 0.05

**Table S4. Impact of the inoculation treatments on the composition of the soil bacterial community.** Identification of significantly affected OTUs between soil microcosms inoculated with Pseudomonas sp. ADP3 (ADP) or the consortium (CONS) as biocomposites (S) or free-living cells (NS), as determined by a generalized linear mixed model (FDR adjusted p-value ≤ 0.05) at **A**) sampling time T1 and **B**) sampling time T2. The soil microcosms were exposed to different atrazine contamination scenarios: treated with atrazine 15 days before inoculation (D-15), treated the same day of inoculation (D0), or not treated (NT).

**A**

| **Sampling time T1** | **Number of significant OTUs** | **% of significant OTUs** |
| --- | --- | --- |
| **D-15** | **80** | **19.41** |
| ADP_NS vs ADP_S | 10 | 2.42 |
| ADP_NS vs NI_NS | 9 | 2.18 |
| ADP_S vs NI_S | 4 | 0.97 |
| CONS_NS vs CONS_S | 23 | 5.58 |
| CONS_NS vs NI_NS | 6 | 1.45 |
| CONS_S vs NI_S | 19 | 4.61 |
| CONS_NS vs ADP_NS | 14 | 3.39 |
| ADP_S vs CONS_S | 40 | 9.70 |
| NI_NS vs NI_S | 7 | 1.69 |
| **D0** | **77** | **18.68** |
| ADP_NS vs ADP_S | 4 | 0.97 |
| ADP_NS vs NI_NS | 14 | 3.39 |
| ADP_S vs NI_S | 20 | 4.85 |
| CONS_NS vs CONS_S | 1 | 0.24 |
| CONS_NS vs NI_NS | 9 | 2.18 |
| CONS_S vs NI_S | 1 | 0.24 |
| CONS_NS vs ADP_NS | 7 | 1.69 |
| ADP_S vs CONS_S | 32 | 7.75 |
| NI_NS vs NI_S | 3 | 0.72 |
| **NT** | **38** | **9.22** |
| ADP_NS vs ADP_S | 2 | 0.48 |
| ADP_NS vs NI_NS | 4 | 0.97 |
| ADP_S vs NI_S | 2 | 0.48 |
| CONS_NS vs CONS_S | 1 | 0.24 |
| CONS_NS vs NI_NS | 2 | 0.48 |
| CONS_S vs NI_S | 12 | 2.91 |
| CONS_NS vs ADP_NS | 0 | 0 |
| ADP_S vs CONS_S | 10 | 2.42 |
| NI_NS vs NI_S | 15 | 3.64 |

**B**

| **Sampling time T2** | **Number of significant OTUs** | **% of significant OTUs** |
| --- | --- | --- |
| **D-15** | **74** | **17.96** |
| ADP_NS vs ADP_S | 16 | 3.88 |
| ADP_NS vs NI_NS | 4 | 0.97 |
| ADP_S vs NI_S | 10 | 2.42 |
| CONS_NS vs CONS_S | 4 | 0.97 |
| CONS_NS vs NI_NS | 10 | 2.42 |
| CONS_S vs NI_S | 9 | 2.18 |
| CONS_NS vs ADP_NS | 3 | 0.72 |
| ADP_S vs CONS_S | 34 | 8.25 |
| NI_NS vs NI_S | 6 | 1.45 |
| **D0** | **227** | **55.09** |
| ADP_NS vs ADP_S | 73 | 17.71 |
| ADP_NS vs NI_NS | 5 | 1.21 |
| ADP_S vs NI_S | 9 | 2.18 |
| CONS_NS vs CONS_S | 4 | 0.97 |
| CONS_NS vs NI_NS | 107 | 25.97 |
| CONS_S vs NI_S | 21 | 5.09 |
| CONS_NS vs ADP_NS | 123 | 29.85 |
| ADP_S vs CONS_S | 60 | 14.56 |
| NI_NS vs NI_S | 58 | 14.07 |
| **NT** | **36** | **8.37** |
| ADP_NS vs ADP_S | 0 | 0 |
| ADP_NS vs NI_NS | 3 | 0.72 |
| ADP_S vs NI_S | 1 | 0.24 |
| CONS_NS vs CONS_S | 11 | 2.66 |
| CONS_NS vs NI_NS | 3 | 0.72 |
| CONS_S vs NI_S | 6 | 1.45 |
| CONS_NS vs ADP_NS | 0 | 0 |
| ADP_S vs CONS_S | 12 | 2.91 |
| NI_NS vs NI_S | 0 | 0 |

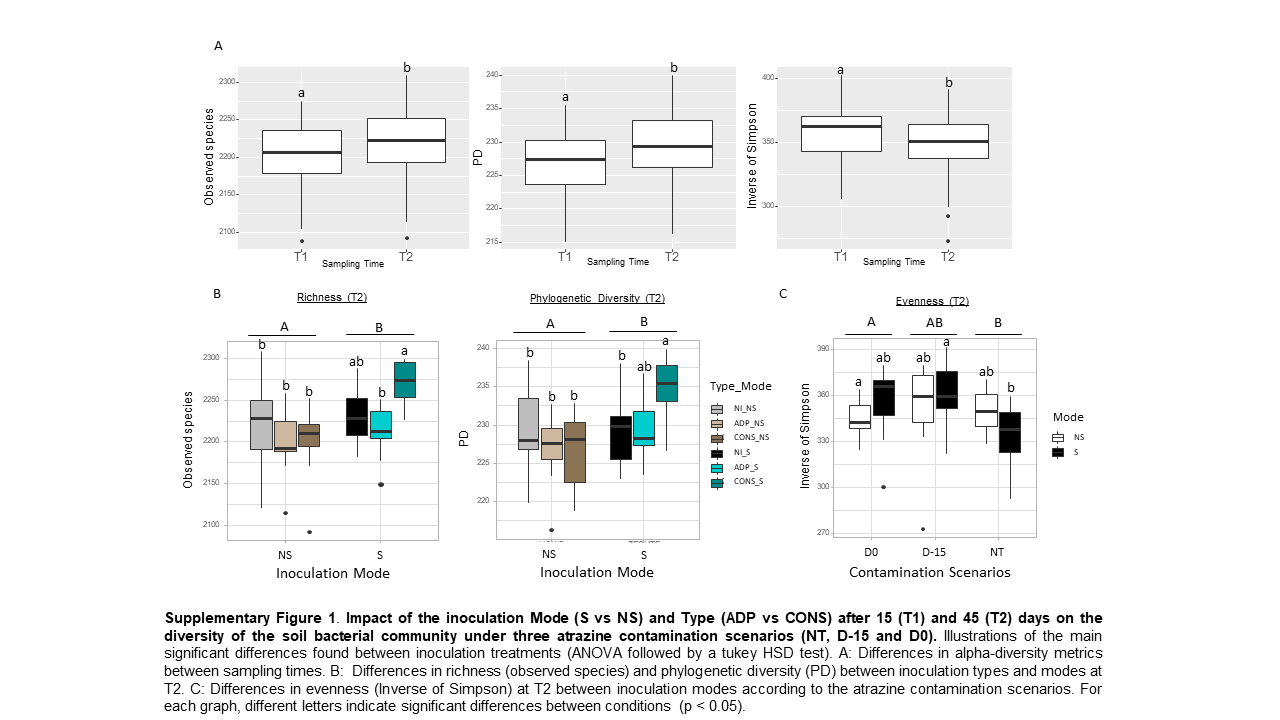

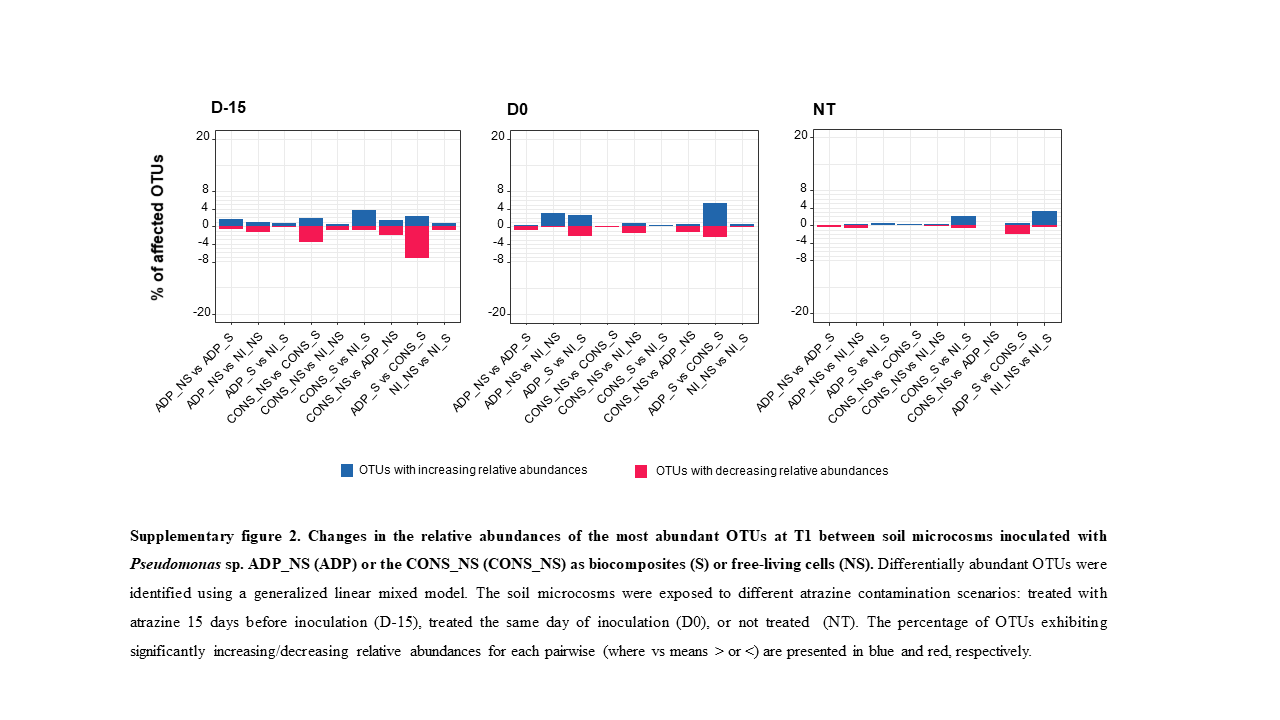
